# Highly amine-reactive graphene-oxide EM grids for biochemical surface modification in aqueous buffer

**DOI:** 10.1101/2023.11.08.566175

**Authors:** Simon H.J. Brown, James C. Bouwer, Scott B. Cohen

## Abstract

Graphene oxide (GO), an oxidized derivative of graphene, has found application in cryo-electron microscopy (cryo-EM) as a hydrophilic and transparent solid support on which to adsorb biological macromolecules, providing an alternative to traditional aqueous films. Current applications generally adsorb the macromolecule directly onto unmodified GO or modify the GO surface with polyethylene glycol-amine reagents. This nucleophilic amine reaction must be performed in an aprotic organic solvent and therefore precludes the use of biological samples such as nucleic acids and peptides. The utility of GO could be expanded by the ability to covalently modify its surface with biochemical affinity reagents such as small- molecule metabolites, peptides, or nucleic acids, in aqueous buffer at neutral pH. Presented here is a chemical procedure that converts all oxygen functionalities of GO to highly amine- reactive glycidyl epoxide groups, achieved without the need of specialized laboratory equipment. We show that single sheets of glycidyl epoxide-modified GO react on the EM grid with primary amines at micromolar concentrations in minutes at room temperature in aqueous buffer. Given the ease of derivatizing biochemical reagents with amines, the chemistry described here will enable imaging of macromolecules immobilized on GO through specific biochemical and biologically relevant binding interactions.

## 1. Introduction

Technologies underlying macromolecular structure determination with single-particle cryogenic electron microscopy (cryo-EM) continue to progress at a rapid pace (*1-4*). Automated acquisition platforms combined with the development of ever-faster direct electron detectors (*5,6*) and the increasing ability to handle Terabytes of data have enabled data collection at hundreds of images per hour and data sets of tens of thousands of images and millions of particles. Furthermore, the researcher has a variety of processing softwares that can generate results in a matter of hours (*7-10*). However, the stringent requirements of a high-quality cryo-EM specimen – thin films of vitrified ice with monodisperse particles presented with a broad distribution of orientations – continues to be an empirical experiment and the major bottleneck in the overall cryo-EM workflow. Common challenges include aggregation of the macromolecule, denaturation, or preferred orientation; these undesirable outcomes have been attributed to the macromolecule encountering the air-water interface through diffusion in the thin aqueous film prior to vitrification, estimated to occur on the millisecond time scale (*11-14*).

An alternative approach to cryo-EM specimen preparation receiving increasing attention is the application of a layer of graphene-oxide (GO) (**Fig. 1**) over the pores of the grid; the macromolecule is adsorbed to the GO, preventing diffusion to the air-water interface (*15*). Compared to other carbonaceous materials such as amorphous carbon, GO is virtually transparent under the electron beam, thereby maintaining the contrast of traditional thin aqueous films; GO is also hydrophilic, structurally rigid, and inexpensive. High-resolution cryo-EM

**Figure 1.**
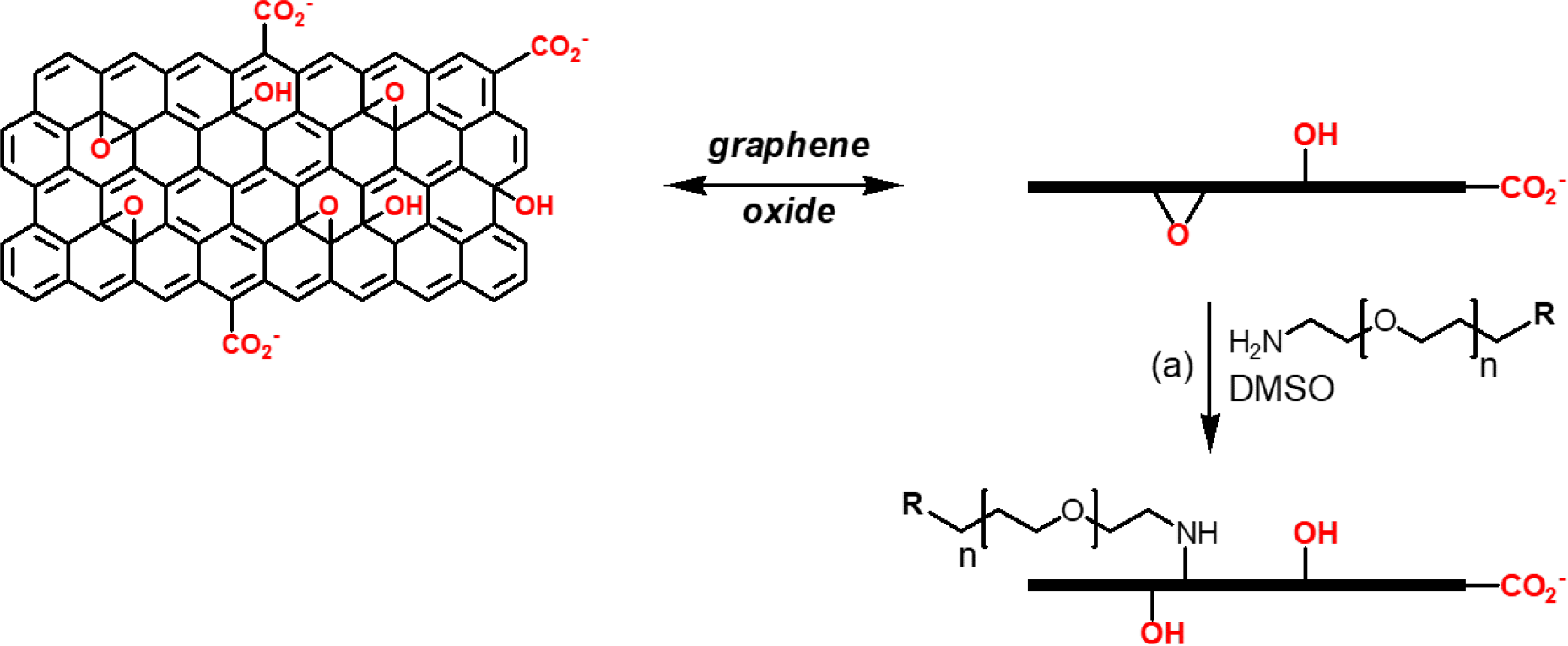
Structure of a sheet of graphene-oxide (GO). GO preparations are heterogeneous with respect to sheet size and extent of oxidation. PEG-amines have been used to modify the surface properties of GO through nucleophilic addition of primary amines to epoxide groups in aprotic organic solvents (*20,21*).

structures of lysenin (*16*) and a separase-securin complex (*17*) on GO demonstrated its ability to circumvent the aggregation or preferred orientation, respectively, that these molecules displayed in aqueous films.

As the oxidized form of graphene, sheets of GO contain three different oxygen functional groups: epoxide, phenol, and carboxylic acids located around the edges of the sheet (**Fig. 1**). The epoxides of GO are reactive towards primary amines in polar aprotic organic solvents such as dimethylformamide or dimethyl sulfoxide (DMSO); the aromatic nature of GO necessitates millimolar concentrations of amine and either heat or extended reaction times (12-24 hr) for this nucleophilic addition to proceed (*18,19*). This reaction has been performed on GO-coated grids with PEGylated amines (**Fig. 1**) to soften the GO surface, resulting in superior particle retention and distribution (*20*). In an analogous manner, applying an amine-modified substrate of the covalent ‘SpyCatcher’ coupling system enabled specific covalent coupling with the complementary Spy-tagged protein to the GO surface (*21*).

We desired a more generalizable means to functionalize the GO surface that would not require protein affinity tags and enable direct coupling with biochemically relevant substrates such as small-molecule metabolites, peptides, or nucleic acids. Covalent coupling of these biochemical substrates to GO would inherently require transition away from organic solvents into an aqueous buffer system; furthermore, the chemical reactivity of such a system would have to be increased by several orders of magnitude to accommodate the typical micromolar concentrations of biochemistry. Here we show that appending glycidyl epoxide groups to GO satisfies these requirements, yielding a highly amine-reactive GO-grid surface. Amine chemistry was chosen given the breadth and ease of derivatizing biochemical reagents with primary amines. We present a simple and inexpensive chemical procedure that converts essentially all oxygen functionalities of the GO to glycidyl epoxides. Applying gold nanoparticle-amine conjugates in aqueous buffer reveals high-density covalent modification of the GO grid surface in minutes with micromolar amine concentrations at room temperature. The chemistry described should enable “on-grid” biochemistry – imaging of macromolecules immobilized on GO through specific biochemical and biologically relevant binding interactions.

### 2. Chemical modification of GO-coated grids

Our motivation for expanding GO chemistry stems from our interest in nucleic acid polymerases and helicases, wherein the grid surface could be functionalized with synthetic DNA or RNA substrates. We considered nucleophilic amine chemistry the most appropriate and accessible for biochemical systems in aqueous buffer; for example, appending a primary amine to synthetic nucleic acids is standard chemistry in any commercial synthesis service. Amine-based bioconjugate chemistry is often performed with *N*-hydroxy succinimide (NHS) esters as the electrophilic component. However, NHS esters hydrolyse rapidly [t_1/2_ ≤ 5 min at pH 8, 23 °C] and therefore require millimolar concentrations of amine to yield efficient coupling. In contrast, glycidyl epoxides display greater stability in aqueous buffer and high reactivity towards amines, as described below.

To coat holey carbon grids with GO in a controlled and predictable manner we applied the method of Palovcak et al. (*22*), wherein a film of GO on water is lowered onto the grid (**Fig. S1**, see Materials & Methods). GO-coated grids were sufficiently robust to be taken through the following chemical transformations. To convert epoxides and carboxylic acid groups of GO to hydroxyl groups, GO-coated grids were treated with 4-amino-1-butanol in DMSO at 50 °C (**Fig. 2**). This provided a mixture of -OH groups derived from GO and amino- 1-butanol.

**Figure 2.**
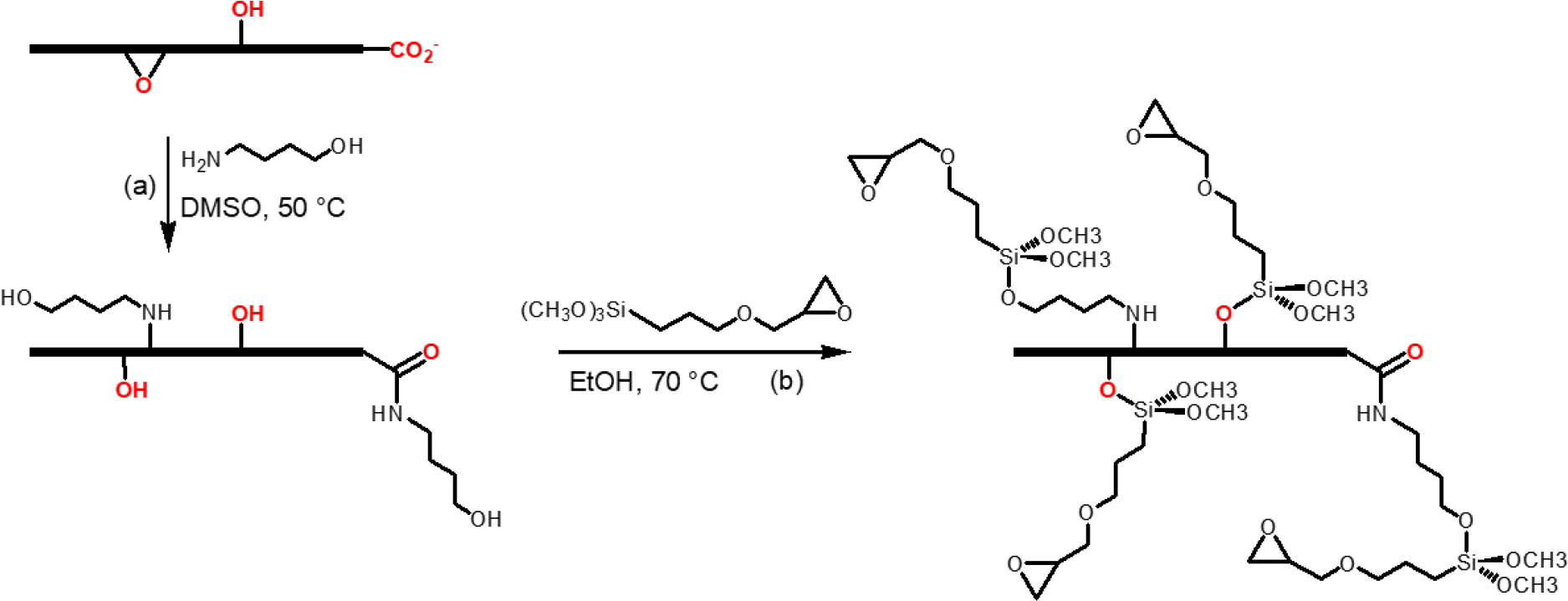
Two-step chemical sequence to produce amine-reactive GO surface.(a) 4-amino-butanol (10 mM), DMSO, 50 °C, 1 hr.(b) 3-glycidoxypropyltrimethoxysilane (50 mM), EtOH, 70 °C, 4 hr.

Silanization is an established method of modifying polyhydroxy molecules and surfaces, and silanization of GO has been reported (*23,24*). The reagent 3- glycidoxypropyltrimethoxysilane couples the oxygen reactivity of a trimethoxysilane with the amine reactivity of a glycidyl epoxide (*25*). Following reaction of GO-coated grids with 4- amino-1-butanol, the grids are treated with a solution of 3-glycidoxypropyltrimethoxysilane in absolute ethanol at 70 °C (**Fig. 2**). This two-step chemical sequence converts essentially all oxygen functionalities of GO to glycidyl epoxides. The cost of these two reagents is negligible compared to the typical cost of holey-carbon grids, and performing the reactions requires only a heat-block and standard 1.5-mL microcentrifuge tubes (**Fig. S1**).

Glycidyl-epoxy modified GO grids were assessed for their transparency and the integrity of the GO crystalline matrix. At moderate underfocus (defocus) levels used for imaging, GO films are virtually transparent; however, sheets of GO can be visualized at very high underfocus. **Figure 3a** shows a single sheet of glycidyl-epoxy GO (approximately 20 μm across) at -10 mm (left) and -100 μm (right) underfocus in low-magnification (LM) mode, illustrating the transparency of the chemically modified grids. GO displays a hexagonal diffraction pattern representing periodicity of the six-carbon lattice (*15,26*); the diffraction displayed by glycidyl-epoxy GO was indistinguishable from unmodified GO (**Fig. 3b**).

**Figure 3.**
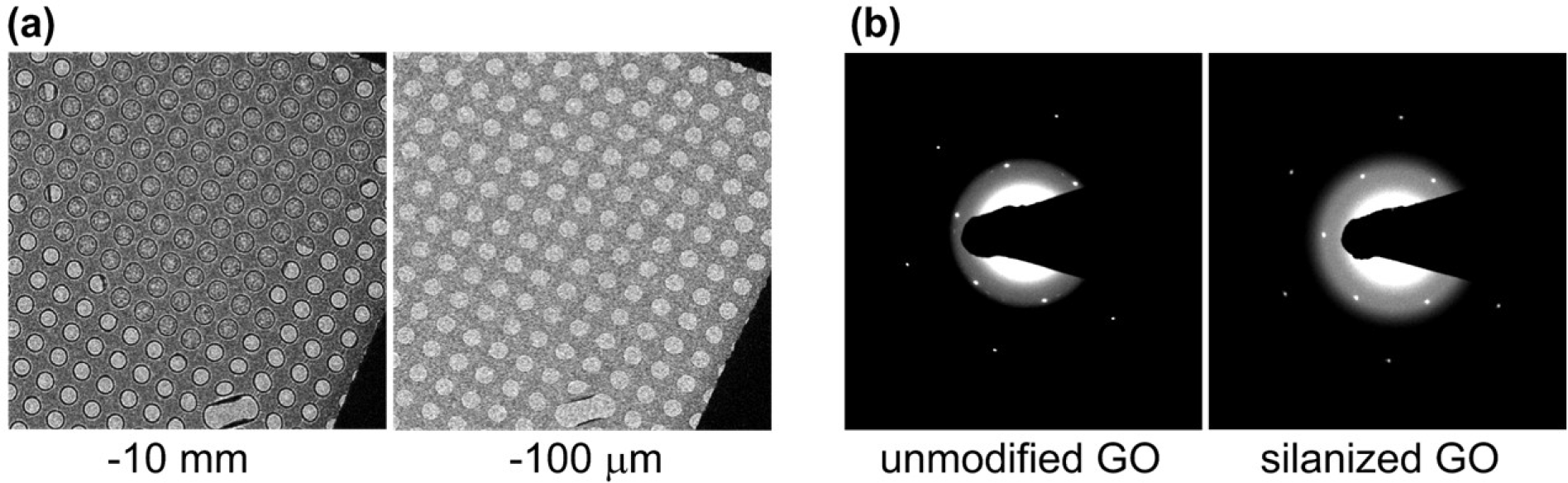
Transparency and integrity of modified GO. (**a**) A single sheet of glycidyl-epoxy modified GO was visualized at high underfocus (left); imaging at low underfocus illustrates the transparent nature of GO. (**b**) A holey-carbon EM grid coated with a single sheet of GO was imaged in diffraction mode, yielding a hexagonal diffraction pattern characteristic of the GO lattice with high periodicity (*15,26*). Diffraction quality after chemical modification was indistinguishable from unmodified GO.

### 3. Amine reactivity of glycidyl-epoxy GO grids

To assess the amine reactivity of glycidyl-epoxy GO grids, a 5-nm gold nanoparticle conjugated to a primary amine was applied (**Fig. 4**). The primary alkyl amine linkage is reflective of typical bioconjugate linkers, such as amine-modified nucleic acids, while the gold nanoparticle provides a visual readout of covalent coupling. A solution at 10 μM amine in HEPES-KOH buffer (pH 8) was applied to a series of grids and incubated at 23 °C; at designated times the reaction was quenched by washing the grid in a beaker of water. Imaging (*Talos Arctica*, 200 kV) showed a time-dependent increase in nanoparticle density (**Fig. 5a-d**), with significant covalent coupling on the time-scale of minutes and largely complete within 30 minutes. To confirm specificity for amine reactivity an analogous 5-nm gold nanoparticle conjugated to a primary hydroxyl (10 μM) was incubated with a glycidyl-epoxy GO grid for 1 hr, revealing sparse non-specifically adsorbed nanoparticles (**Fig. 5f**, red), possibly attributed to the hydrophobic nature of gold nanoparticles.

**Figure 4.**
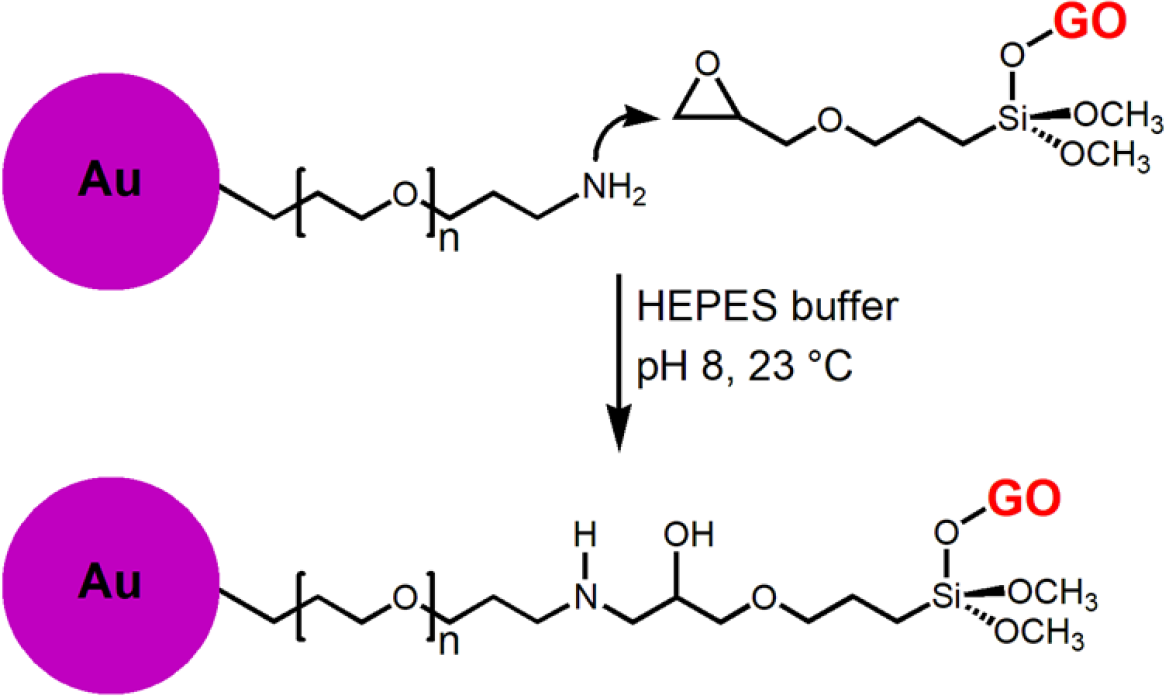
Chemical reaction to quantitatively and visually assess reactivity of terminal glycidyl epoxide with primary amines on GO. Gold nanoparticles (∼5 nm diameter) were covalently modified with a PEG linker (n ∼ 25) bearing a terminal alkyl amine. The coupling was performed in aqueous HEPES-KOH buffer (pH 8) at rt and followed by EM (see **Figure 5**).

**Figure 5.**
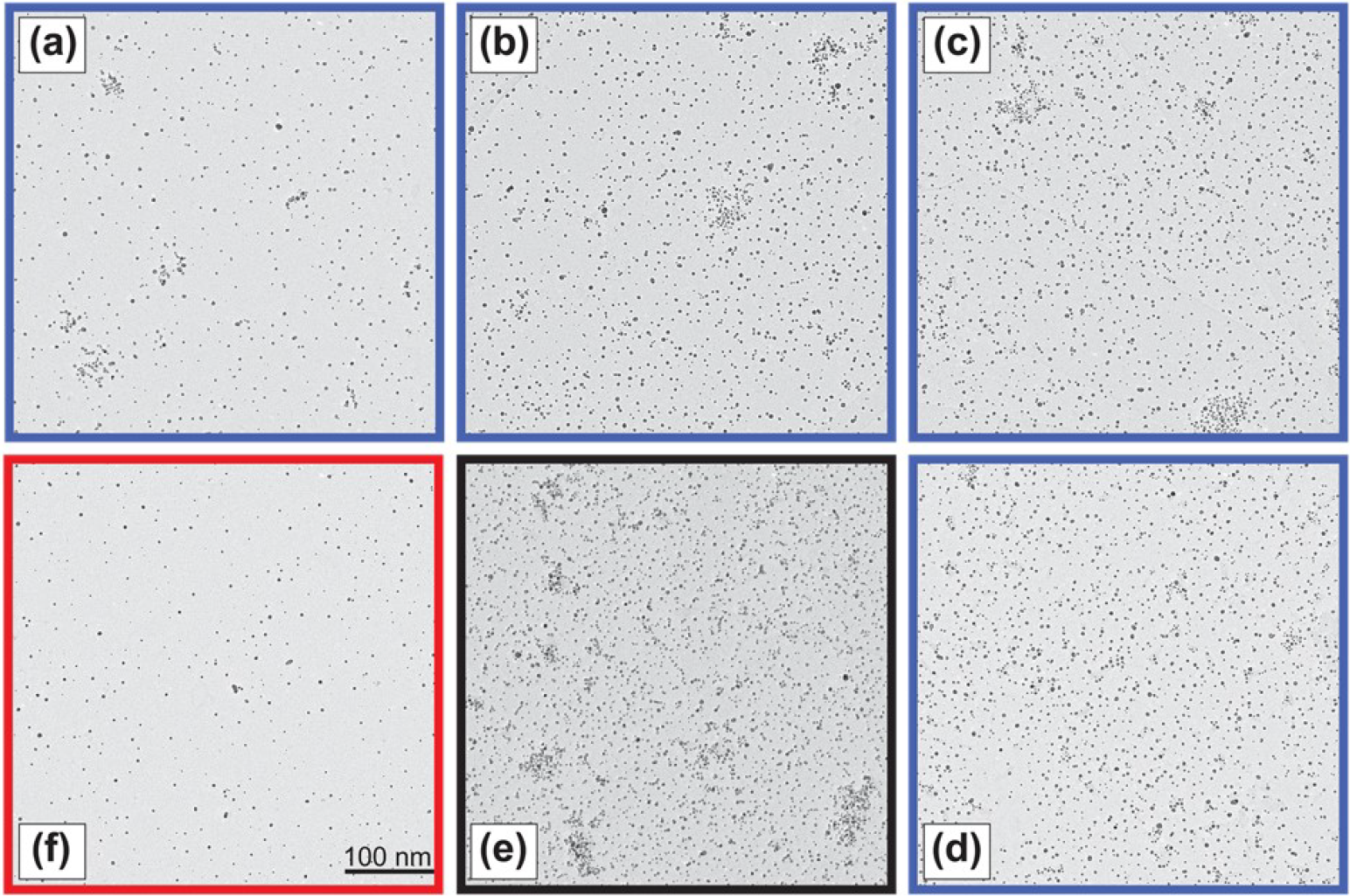
Kinetics of covalent coupling of amine-conjugated gold nanoparticle, visualized by electron microscopy. A modified GO grid was treated with the Au- amine conjugate at a concentration of 10 μM amine in aqueous HEPES-KOH buffer (pH 8) at rt for: (**a**) 5 min; (**b**) 15 min; (**c**) 30 min; (**d**) 60 min. (**e**) Hydrolysis control: the grid was incubated in aqueous HEPES-KOH buffer (pH 8) for 60 min before reaction with Au-amine (10 μM for 60 min) to assess hydrolysis of the glycidyl epoxide; Au particle density was indistinguishable from (**d**), indicating that background hydrolysis is negligible. (**f**) Specificity control: non-specific adsorption of the gold nanoparticle to the grid surface was assessed by incubating the grid with a gold nanoparticle bearing a terminal hydroxyl (-OH) in place of the amine, 10 μM for 60 min. Black scale bar, 100 nm.

Having demonstrated efficient amine reactivity, we assessed the stability of the glycidyl epoxides against background hydrolysis in aqueous buffer. A glycidyl-epoxy GO grid was incubated with HEPES-KOH buffer at 23 °C for 1 hr; the buffer was blotted away and the 10 μM amine-gold nanoparticle reagent was applied. After 1 hr the density of covalent coupling (**Fig. 5e**, black) was indistinguishable from standard treatments (**Fig. 5d**, blue). This analysis indicates that the background hydrolysis reaction of glycidyl epoxides is exceptionally slow, and that amine concentrations significantly lower than 10 μM can still result in high-density covalent coupling on the grid, albeit with longer reaction time. These results were replicated with glycidyl epoxide GO grids that had been stored for 3 months at ambient conditions, further demonstrating the robust nature of these grids.

## 4. Summary

Presented here is chemistry that will expand the utility of graphene-oxide as a support film for single-particle cryo-EM. We developed a two-step chemical procedure that renders the GO surface highly reactive to amines in aqueous buffer at neutral pH. While the high reactivity of the glycidyl-epoxide GO grids is ideally suited to biochemical reagents for which quantities and/or concentrations may be limiting, one could envision the application of reagents to confer more general properties to the grid surface. For example, treating the grids with 4-aminobutryic acid would provide a negatively-charged surface; reaction with *α*-amino acid methyl esters is another route to providing a surface with a wide range of chemical properties.

Our interest has been in the application of nucleic acid substrates, covalently immobilizing these to the GO surface to promote capture of the cognate polymerase or helicase. Compared to the established method of modifying GO with PEGylated amines in organic solvent (*20,21*) the amine-coupling chemistry presented here: (i) occurs in aqueous buffer at neutral pH, thereby greatly expanding the molecular diversity of reagents that can be applied; (ii) is achieved with low-micromolar concentrations of amine, ∼10^2^-10^3^ lower than the millimolar concentrations required for amine addition to the GO-epoxides in organic solvent; (iii) functionalizes essentially all oxygen atoms on the GO, enabling modification at higher density.

## 5. Materials & Methods

This work was performed on *Quantifoil* copper R1.2/1.3 200-mesh grids. Graphene oxide was purchased from *Sigma-Aldrich* (now *Merck*) as a 2 mg/mL dispersion in water (*Merck* #763705); two bottles obtained from different lots, displaying slightly different viscosity and color, were used (see below). 4-Amino-1-butanol (#178330) and 3- glycidoxypropyltrimethoxysilane (#440167) were obtained from *Merck*. Gold nanoparticle- PEG-amine conjugate was custom-synthesized by *NanoPartz* (Loveland CO) and provided as an aqueous suspension at 4 μM nanoparticle.

### Coating grids with graphene-oxide (GO)

We followed the method of Palovcak et al. (*22*), wherein a film of GO on water is lowered onto the grid (**Fig. S1a**). Ten to fifteen grids per batch were glow-discharged in a *PELCO EasiGlow* glow-discharge system at settings of 0.4 mBarr and 15 mAmp for 45 sec. A screen mesh was placed in a Pyrex dish of 5-cm diameter and submerged with water; the water level was ∼1 cm above the mesh. Grids were placed carbon-side up on the mesh, and an inlet tube connected to a peristaltic pump was lowered into the water (**Fig. S1a**). Commercial GO preparations are heterogeneous with respect to the size of the individual sheets and the extent of oxidation, leading to differences in viscosity and color and potentially differences in experimental outcomes, as described below. To prepare a dilute dispersion of GO the procedure of Palovcak et al. uses a dispersant of 5:1 v/v CH_3_OH:H_2_O (*22*). Our aqueous 2 mg/mL GO dispersion (50 μL) was diluted into 950 μL dispersant and mixed with vortexing to provide GO at 100 μg/mL. For one of the *Sigma* preparations this procedure provided an essentially clear homogenous dispersion; however, the other *Sigma* preparation resulted in a suspension with visible particles (aggregates) of GO. We found through empirical trial that an additional 100 μL H_2_O resulted in a clear homogeneous dispersion (GO now at 90 μg/mL). Using a pipette, 1-2 μL drops of GO dispersion were gently applied to the water surface, moving around the entire surface, applying a total of 12 μg GO.

The water was then removed by the peristaltic pump over a period of ∼15 min, lowering the GO layer onto the grids. The GO-coated grids were air-dried on the mesh overnight, transferred to filter paper (GO side up) and air-dried for another 24 hr.

### Reaction of GO with 4-aminobutanol (Fig. 2)

DMSO was stored over 4 Å molecular sieves to minimize water content. A solution of 4-aminobutanol (10 mM) in DMSO was prepared by combining 9.99 mL DMSO and 9.2 μL neat 4-aminobutanol. Into each of ten 1.5-mL microcentrifuge tubes was placed 200 μL of the 10-mM amine solution. A GO-coated grid was placed into each tube, submerged in the reaction solution (**Fig S1b**). The tubes were incubated in a heat block at 50 °C for 1 hr. To retrieve the grid, the entire contents of the tube was poured onto a bed of paper towels. The grid was rinsed in a beaker (∼200 mL) of absolute ethanol for 30 sec, placed GO-side up on filter paper, and air-dried overnight.

### Silanization with 3-glycidoxypropyltrimethoxysilane (Fig. 2)

Safety: this reagent is acutely damaging to eyes – wear safety glasses! The 100-mL bottle comes sealed with a rubber septum; we use a 1-mL syringe equipped with a 25-gauge needle to transfer ∼200 μL into a 1.5-mL microcentrifuge tube, thereby minimising exposure. A solution of 3- glycidoxypropyletrimethoxysilane (50 mM) in absolute ethanol was prepared by combining 9.89 mL ethanol and 110 μL neat silane. Into each of ten 1.5-mL microcentrifuge tubes was placed 200 μL of the 50-mM silane solution. An amine-treated grid from above was placed into each tube, submerged in the reaction solution. The tubes were incubated in a heat block at 70 °C for 4 hr. To retrieve the grid, the entire contents of the tube was poured onto a bed of paper towels. The grid was rinsed in a beaker (∼200 mL) of absolute ethanol for 30 sec, placed GO-side up on filter paper, and air-dried overnight.

### Transparency and diffraction of glycidyl-epoxy modified GO grids (Fig. 3)

A modified grid was loaded into a *Tecnai* T12 transmission electron microscope operating at 120 kV equipped with a *Gatan* Rio4 camera. The grid was scanned at low magnification (∼80-250×) at a defocus of -10 mm, allowing visualization of individual sheets of GO. Having identified a sheet of GO, the magnification was increased to 6800× and centered on an individual pore. The microscope was changed to diffraction mode (diffraction length = 120 mm; C2 set to produce parallel illumination) revealing the hexagonal diffraction.

### Reaction of glycidyl-epoxy modified GO grids with gold nanoparticle-amine conjugate (Figs. 4 & 5)

The 5-nm gold nanoparticle-PEG-amine conjugate was provided as a deep purple aqueous suspension at 4 μM nanoparticle. Nanoparticles were covalently modified with ∼2 molecules of PEG-amine per nm^2^, corresponding to ∼157 amines per nanoparticle and ∼628 μM total amine concentration. A 10-μL aliquot of the nanoparticle suspension was diluted with 620 μL [20 mM HEPES-KOH (pH 8) + 100 mM NaCl] to provide 10 μM amine (630 μL). The light purple solution was sonicated (*Shesto* model UT8031/EUK, 100 W) at high power for 1 min. A set of four glycidyl-epoxy GO grids were placed on a glass plate; to each was added 8 μL of the 10-μM amine-nanoparticle solution and the reactions incubated at 23 °C. At reaction times of 5, 15, 30, and 60 min a grid was rinsed in a beaker (∼200 mL) of Milli-Q H_2_O for 30 sec to quench the reaction. Residual H_2_O was gently side-blotted away with filter paper, and the grids were air-dried on filter paper overnight. Grids were imaged dry on a *Talos Arctica*, operating at 200 kV, equipped with a Flacon-III direct electron detector, acquiring at a pixel size of 1.18 Å.

## Acknowledgement

The authors thank Eugene Palovcak and Yifan Cheng (Department of Biochemistry & Biophysics, UC San Francisco) for generously sharing their graphene-oxide methods before publication. SBC acknowledges support from the Ernest & Piroska Major Foundation, Cancer Council New South Wales, and Kids Cancer Alliance (Australia).

---

**Figure S1.**
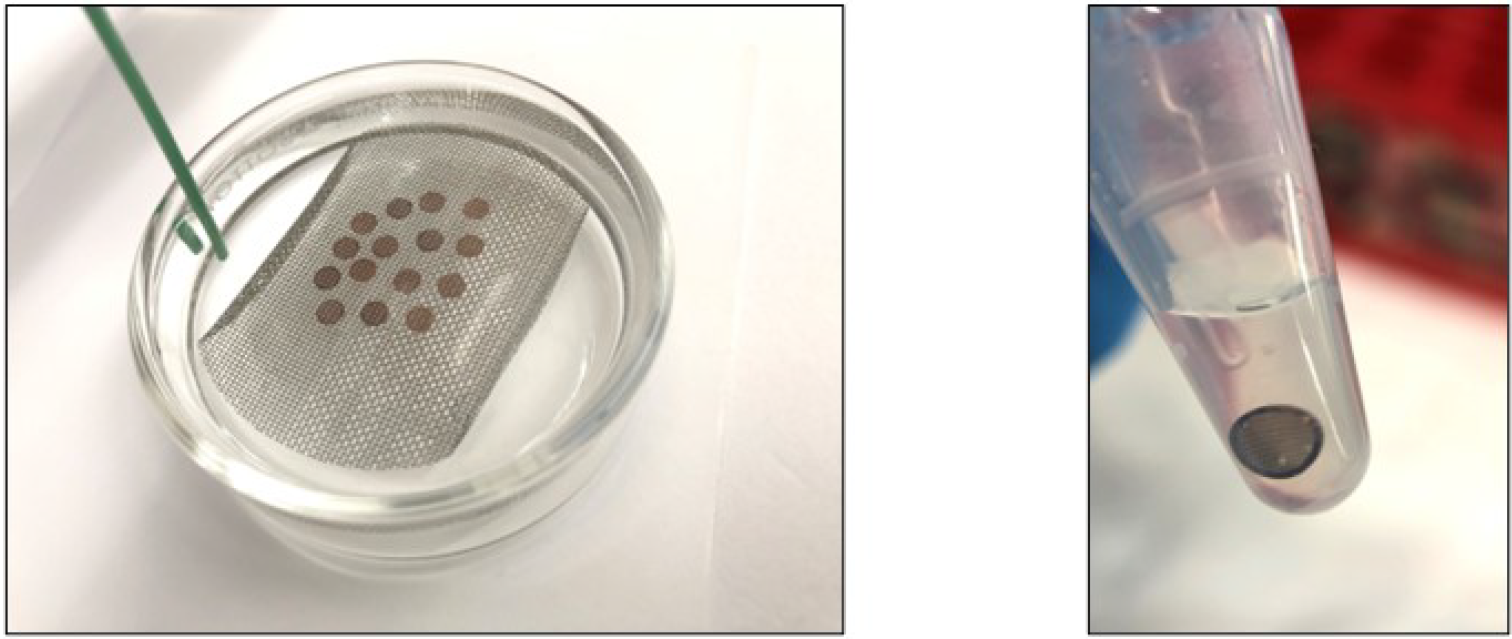
(**a**) Laboratory set-up for coating holey-carbon EM grids with GO (*22*). Grids are submerged in water on a wire mesh. A dispersion of GO has been applied dropwise to the surface of the water forming a monolayer (GO not visible), and the GO is lowered onto the grids by removing the water with a pump (green tubing). (**b**) Chemical modification of grids is carried out in standard 1.5-mL microcentrifuge tubes.

## Notes

### Competing Interest Statement

The authors have declared no competing interest.

